# A multidimensional functional fitness score is a stronger predictor of type 2 diabetes than obesity parameters in cross sectional data

**DOI:** 10.1101/580860

**Authors:** Pramod Patil, Poortata S Lalwani, Harshada B Vidwans, Shubhankar A Kulkarni, Deepika Bais, Manawa M Diwekar-Joshi, Mayur Rasal, Nikhila Bhasme, Mrinmayee Naik, Shweta Batwal, Milind G Watve

**Affiliations:** Indian Institute of Science Education and Research, Pune (IISER – P) India; Deenanath Mageshkar Hospital, Pune India; Department of Psychology, University of Michigan, Ann arbor, MI - 48109, USA

**Author notes:** Corresponding author: Milind G Watve, Indian Institute of Science Education and Research, Pune (IISER – P) India. Dr. Homi Bhabha Rd, Ward No. 8, NCL Colony, Pashan, Pune, Maharashtra 411008, Phone(O), Fax (O).

## Abstract

**Objective:** We examine here whether multidimensional functional fitness is a better predictor of type 2 diabetes as compared to morphometric indices of obesity such as body mass index (BMI) and waist to hip ratio (WHR).

**Research design and method:** We analysed retrospective data of 663 volunteer participants (285 males and 378 females between age 28 and 84), from an exercise clinic in which every participant routinely undergoes a health related physical fitness (HRPF) assessment consisting of 15 different tasks examining 8 different aspects of functional fitness.

**Results:** The odds of being diabetic in the highest quartile of BMI were not significantly higher than that in the lowest quartile in either of the sexes. The odds of being a diabetic in the highest WHR quartile were significantly greater than the lowest quartile in females (OR = 4.54 (1.95, 10.61) as well as in males (OR = 3.81 (1.75, 8.3). In both sexes the odds of being a diabetic were significantly greater in the lowest quartile of HRPF score than the highest (males OR = 10.52 (4.21, 26.13); females OR = 10.50 (3.53, 31.35)). HRPF was not correlated with BMI in both sexes but was negatively correlated with WHR. After removing confounding, the predictive power of HRPF was significantly greater than that of WHR.

**Conclusion:** Multidimensional functional fitness score was a better predictor of type 2 diabetes than obesity parameters in the Indian population.

## Introduction

In the mainstream thinking in the field of type 2 diabetes mellitus (T2DM) obesity is believed to be causal to insulin resistance, and subsequently to T2DM (1). Many shortcomings and paradoxes in this view are becoming apparent with increasing research inputs. The direction of causality between obesity, insulin resistance and hyperinsulinemia is debated. While the classical view presumes obesity to be primary, giving rise to insulin resistance and hyperinsulinemia as a compensatory response of the body, an upcoming view is that hyperinsulinemia has a causal role in obesity (2–5). Downregulation of insulin expression by insulin gene dosage (6), pharmaceutical suppression of insulin (7) or fat cell specific insulin receptor knockouts (8) protect against obesity. Epidemiological evidence is predominantly associative and does not clearly show causal relationship of obesity with insulin resistance and hyperinsulinemia. In any case the association between obesity and insulin resistance is quantitatively quite weak throughout the globe with the variance explained being below 15 % in majority of studies (9). Further, the association varies between populations, normal weight T2DM being more common in south Asians (10–11).

If obesity is a poor predictor of T2DM, one needs to look for alternative and more reliable predictors. There have been arguments and evidence about fitness being more important than fatness (12–19). However, most studies have looked at a single dimension of fitness such as grip strength (16–17), lower body strength (18–19) or cardiorespiratory fitness (12–15). Most of these studies find that some form of functional fitness has a protective role against T2DM and many other life-style related disorders. However, fitness is a multidimensional concept which is difficult to capture in a single task performance test. Whether a multidimensional fitness score has greater predictive power than a single dimension or single task has not been addressed seriously. Currently there is no standardized set of test protocols for assessing multidimensional functional fitness. We found an opportunity to examine multidimensional fitness score as a predictor of T2DM using retrospective data from an exercise clinic in Pune, India which routinely conducted a multidimensional fitness test at entrance for every new participant. We use this opportunity here to make a preliminary assessment of whether it gives us a predictability substantially greater than that given by obesity parameters.

### The clinic and the patient group

Deenanath Mangeshkar Hospital and Research Centre, a multi-speciality hospital in Pune city, India opened an exercise clinic namely BILD (behavioural intervention for lifestyle disorders) clinic in 2017. It serves as an exercise training centre for prevention as well as treatment for neuro-orthopedic, metabolic and endocrine problems. The exercise prescription is personalized depending upon the problem addressed along with the capacity and physical limitations of the individual. In order to record the capacity and limitations of the patients, the centre conducts a routine health related physical fitness (HRPF) examination of every entrant. HRPF is a multidimensional functional fitness test consisting of 15 small tasks that examine different dimensions of physical fitness including abdominal plasticity, balance, endurance, flexibility, nerve-muscle coordination, muscle strength, core strength and agility. The set of tests, described in detail in the supplementary information, gives a differential score to each of the fitness components and a total composite fitness score. During past one year the clinic had performed over 800 HRPF assessments covering age groups between 18 and 84. We noted that type 2 diabetics in the sample ranged between ages 28 and 84 and therefore we used the same range for the non-diabetic counterpart of the sample. This resulted in the selection of 285 males and 378 females out of which 81 males and 69 females were type 2 diabetic.

## Results

BMI did not correlate with age significantly in either of the sexes (males r = −0.136, NS; females r = 0.085, NS). In both sexes WHR correlated positively with age (males r = 0.287, p < .001; females r = 0.227, p<0.001). In males, the HRPF score decreased with age but the rate of decrease was significantly steeper in diabetics (r^2^ = 0.451 and regression slope = −0.961, CI =-0.72 to – 1.28) than non-diabetics (r^2^ = 0.342, slope = −0.625, CI = −0.505 to – 0.74). The pattern was similar in females but the difference in the slope was not significant (diabetics r^2^ = 0.342 and slope = −0.747, CI = −0.531 to −1.03)) (non-diabetics r^2^ = 0.296, slope = −0.642, CI = −0.523 to −0.7451)). Nevertheless the best fit regression line for diabetics lay below that for non-diabetics throughout the age range indicating that at any age group diabetics were less fit than non-diabetics (figure 1).

**Figure 1:**
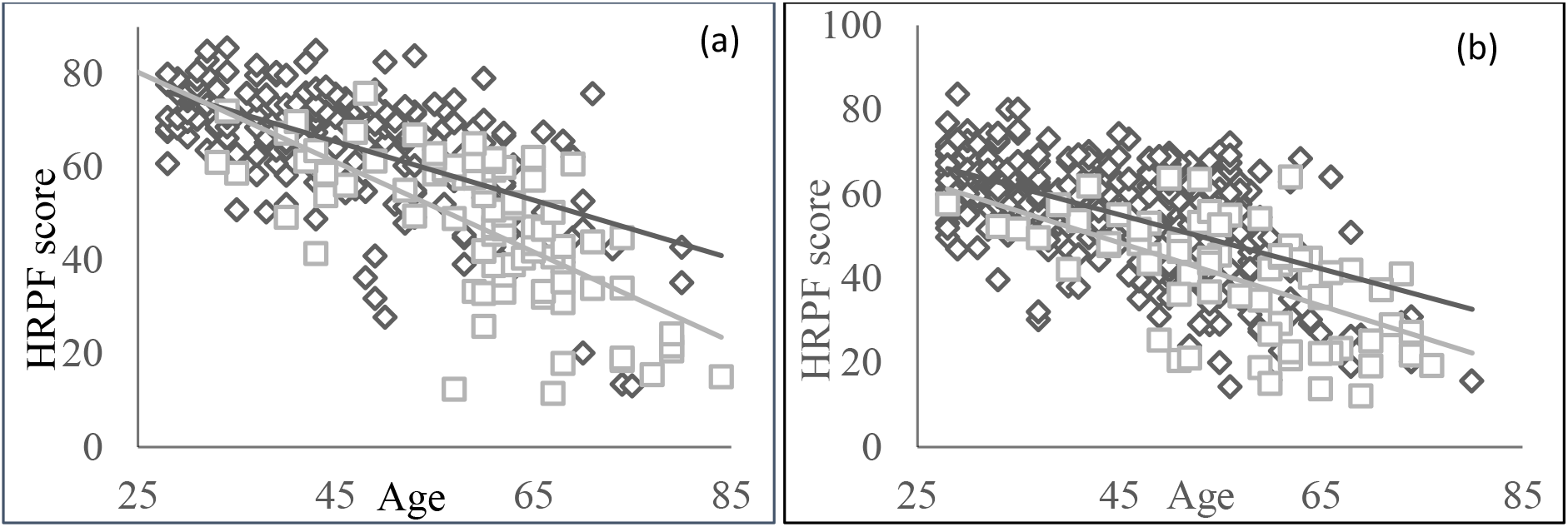
Distribution of diabetics (grey-square) and non-diabetics (black - diamond) with respect to HRPF scores (a) males (b) female

The different components of functional fitness were intercorrelated positively but the *r^2^* values were small, over 95 % of them ranging between 0.00001 and 0.25 with occasional outliers going up to 0.42 (figure 2, table 1). The components were significantly positively correlated to the total score as expected in both sexes and the component that explained maximum variance in HRPF was muscle strength (63.2% in males and 40.3% in females).

**Figure 2:**
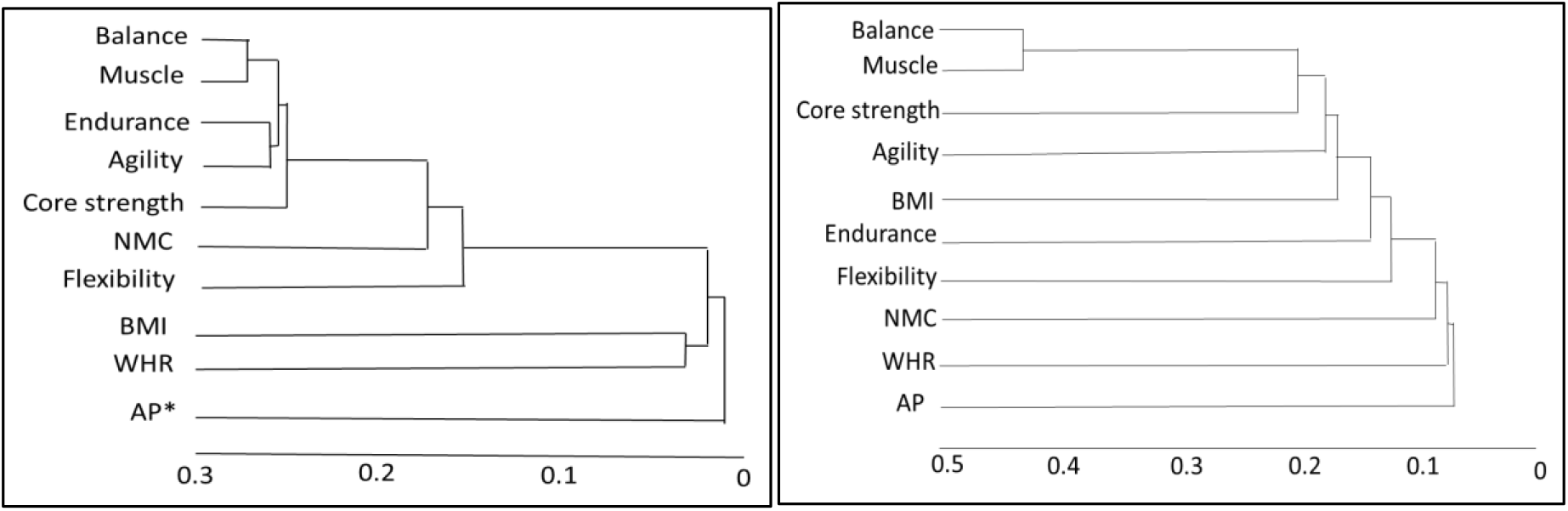
(a) males (b) females: clustering of different components of functional fitness based on the pairwise coefficients of determination, using nearest neighbour clustering. Note that most coefficients are below 0.3 with the exception of balancing and muscle strength in females. Thus no single component represents overall fitness very well.

**Table 1:** Correlation between components of functional fitness and morphometric indices. Significant correlations after applying Bonferroni correction are indicated by bold font. Pair wise (r) values are tabulated in the two sexes. Note that total HRPF score is most consistently correlated with all other measures followed by balance, whereas BMI, WHR and abdominal plasticity are not frequently correlated.

| **Females** | HRPF<br>score | BMI | WHR | AP<br>Score | Balance | Endu | Flex | NMC | Musc | Agil |
| --- | --- | --- | --- | --- | --- | --- | --- | --- | --- | --- |
| BMI | -0.036 |  |  |  |  |  |  |  |  |  |
| WHR | **-0.239** | 0.147 |  |  |  |  |  |  |  |  |
| AP Score | **0.198** | **-0.187** | **-0.271** |  |  |  |  |  |  |  |
| Balance | **0.617** | **-0.430** | **-0.275** | 0.123 |  |  |  |  |  |  |
| Endurance | **0.419** | 0.020 | -0.119 | 0.085 | **0.260** |  |  |  |  |  |
| Flexibility | **0.419** | -0.060 | -0.002 | **0.196** | **0.275** | **0.205** |  |  |  |  |
| NMC | **0.284** | -0.053 | -0.187 | 0.046 | **0.188** | 0.163 | 0.029 |  |  |  |
| Muscle<br>strength | **0.635** | **-0.409** | **-0.170** | 0.162 | **0.655** | **0.274** | **0.349** | **0.170** |  |  |
| Agility | **0.593** | -0.089 | -0.143 | 0.062 | **0.412** | **0.362** | **0.174** | **0.285** | **0.437** |  |
| Core<br>Strength | **0.527** | **-0.247** | -0.122 | 0.110 | **0.442** | **0.276** | **0.215** | **0.169** | **0.449** | **0.344** |
| **Male** | HRPF<br>score | BMI | WHR | AP<br>Score | Balance | Endu | Flex | NMC | Musc | Agil |
| BMI | **-0.213** |  |  |  |  |  |  |  |  |  |
| WHR | **-0.231** | **0.342** |  |  |  |  |  |  |  |  |
| AP Score | **0.347** | -0.076 | 0.049 |  |  |  |  |  |  |  |
| Balance | **0.740** | **-0.323** | **-0.216** | 0.072 |  |  |  |  |  |  |
| Endurance | **0.630** | -0.019 | -0.177 | 0.101 | **0.379** |  |  |  |  |  |
| Flexibility | **0.524** | -0.080 | -0.153 | 0.135 | **0.279** | **0.243** |  |  |  |  |
| NMC | **0.485** | 0.066 | **-0.231** | 0.001 | **0.282** | **0.368** | 0.102 |  |  |  |
| Muscle | **0.795** | **-0.233** | -0.164 | 0.154 | **0.531** | **0.408** | **0.397** | **0.289** |  |  |
| Agility | **0.726** | -0.034 | -0.143 | 0.122 | **0.428** | **0.502** | **0.210** | **0.431** | **0.464** |  |
| Core<br>Strength | **0.666** | -0.189 | -0.136 | 0.072 | **0.441** | **0.404** | **0.299** | **0.313** | **0.502** | **0.401** |

After correcting for age, HRPF score was not correlated with BMI either in diabetics or non-diabetics in both sexes. But BMI was negatively correlated with some of the components of functional fitness namely abdominal plasticity in females (r = −0.187, p <0.001), balance (in males r = −0.323, p < 0.001 and in females −0.430, p < 0.0001), muscle strength (in males r = −0.233, p< 0.001; in females r = −0.409 p <0.001) and core strength in females (r = −0.247, p < 0.001). WHR was significantly negatively correlated with age corrected HRPF (males r = −0.231, p< 0.001; females r = −0.239, p< 0.001). Although statistical significance was seen, the coefficients of determination in both the cases were very small (0.052 and 0.057). BMI was not significantly correlated with WHR in both sexes. Thus neither BMI nor WHR reflected on functional fitness very well.

The odds of finding a diabetic in the lowermost quartile of BMI was significantly greater than that in the highest quartile, contrary to the expectation in males (OR= 0.41 (0.19, 0.89); but the not in females OR = 0.65 (0.31, 1.37)) (figure 3). The odds of being a diabetic in the highest WHR quartile were significantly greater than the lowest quartile in females (OR = 4.54 (1.95, 10.61)) as well as in males (OR = 3.81 (1.75, 8.3)). In both sexes the odds of being a diabetic were significantly greater in the lowest quartile of HRPF score than the highest (males OR = 10.52 (4.21, 26.13); females OR = 10.50 (3.53, 31.35)). In both sexes the ORs for HRPF were over two fold those of the corresponding ORs for WHR but the differences were not statistically significant. However when data on the two sexes were pooled with nested ranking, the odds ratio for HRPF (OR = 9.78 (4.93, 19.8)) was significantly greater than the OR for WHR (4.03 (2.09, 7.08); p one tailed = 0.03). Therefore the predictive power of HRPF appears to be greater than WHR and BMI.

**Figure 3:**
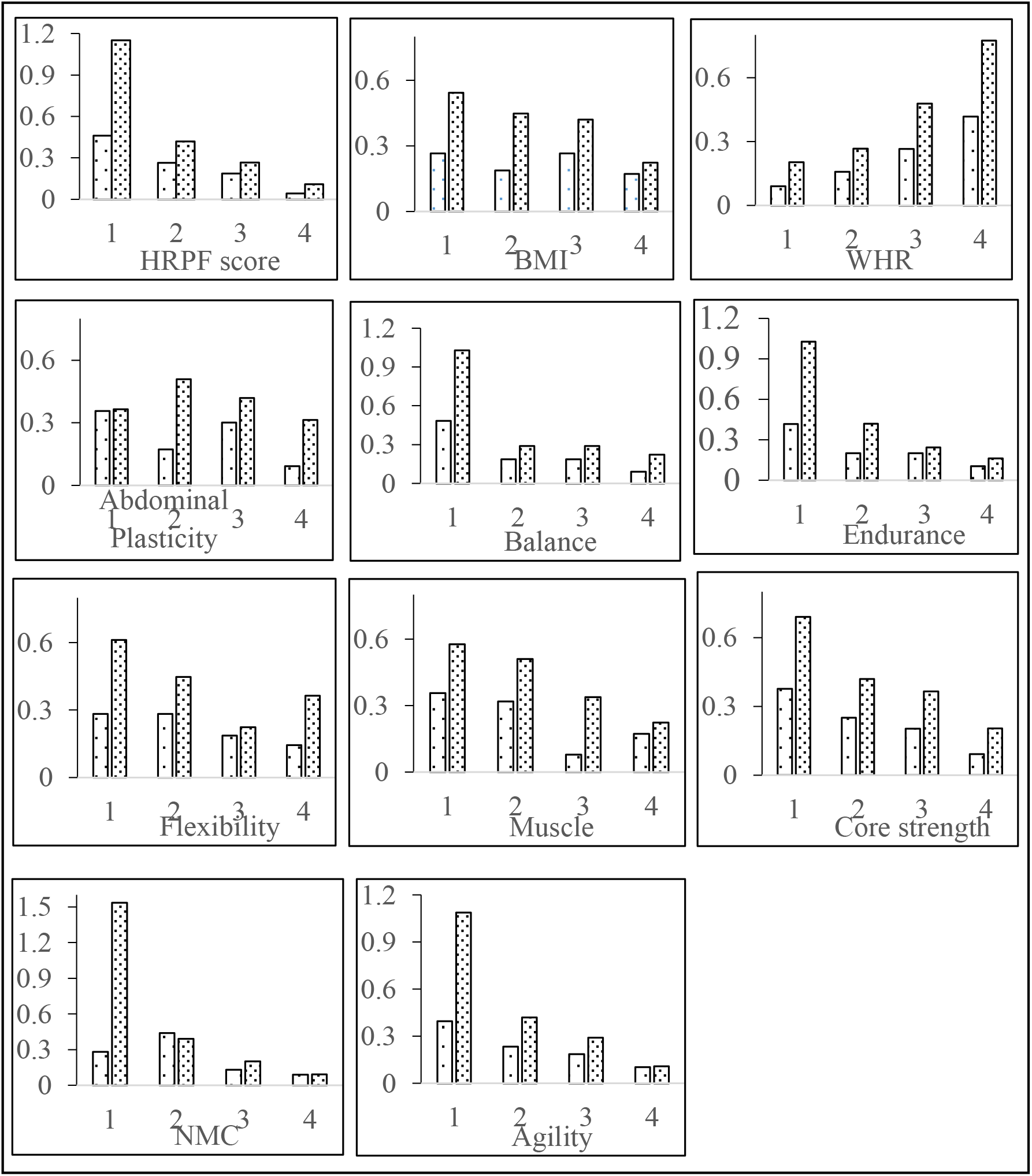
Odds of being Type 2 diabetic among quartiles of different components of functional fitness in comparison with morphometric indices. Males-dense dots, females – sparse dots.

The odds ratios of the lowest to highest quartiles of some of the individual fitness component scores (figure 3) were also significant. However the OR of individual fitness components were lower than the total HRPF score and not significantly different from each other. The only exception was nerve-muscle coordination in males (OR =16.63 (6.36, 35.6)) for which the OR was greater than the OR for HRPF but the difference was not significant. However it was significantly greater than the OR for WHR (p one tailed 0.019). The predictive power of the multidimensional functional fitness generally appears to be better than most of the single task scores or single dimensions of functional fitness.

Since WHR and HRPF were good predictors of diabetes but not BMI, we tested whether WHR and HRPF were independent predictors. After correcting for WHR, HRPF remained a significant predictor although the odds ratio decreased in males (OR = 5.05 (2.33, 10.98)). In contrast the OR for WHR corrected HRPF increased in females (OR = 22.3 (6.59, 75.5)). Reciprocally after correcting for HRPF, WHR remained a significant predictor with a marginal decrease in odds ratios (males 3.766 (1.7, 8.37); females (2.58 (1.18, 5.63)). This indicates that both WHR and HRPF are independent predictors. Moreover after removing the confounding the OR for HRPF in females was significantly greater than OR for WHR (p one tailed 0.0015). Also with both sexes pooled the difference was statistically significant (p one tailed = 0.0068). Thus with or without confounding, multidimensional functional fitness score appears to have a greater predictive value than the obesity parameters BMI and WHR.

In a scatter along the HRPF and WHR coordinates, diabetic and non-diabetic groups formed significantly different clusters (Figure 4) in females (PerMANOVA, F = 30.18, P = 0.001) as well as males (PerMANOVA, F = 42.92, P = 0.001).

**Figure 4:**
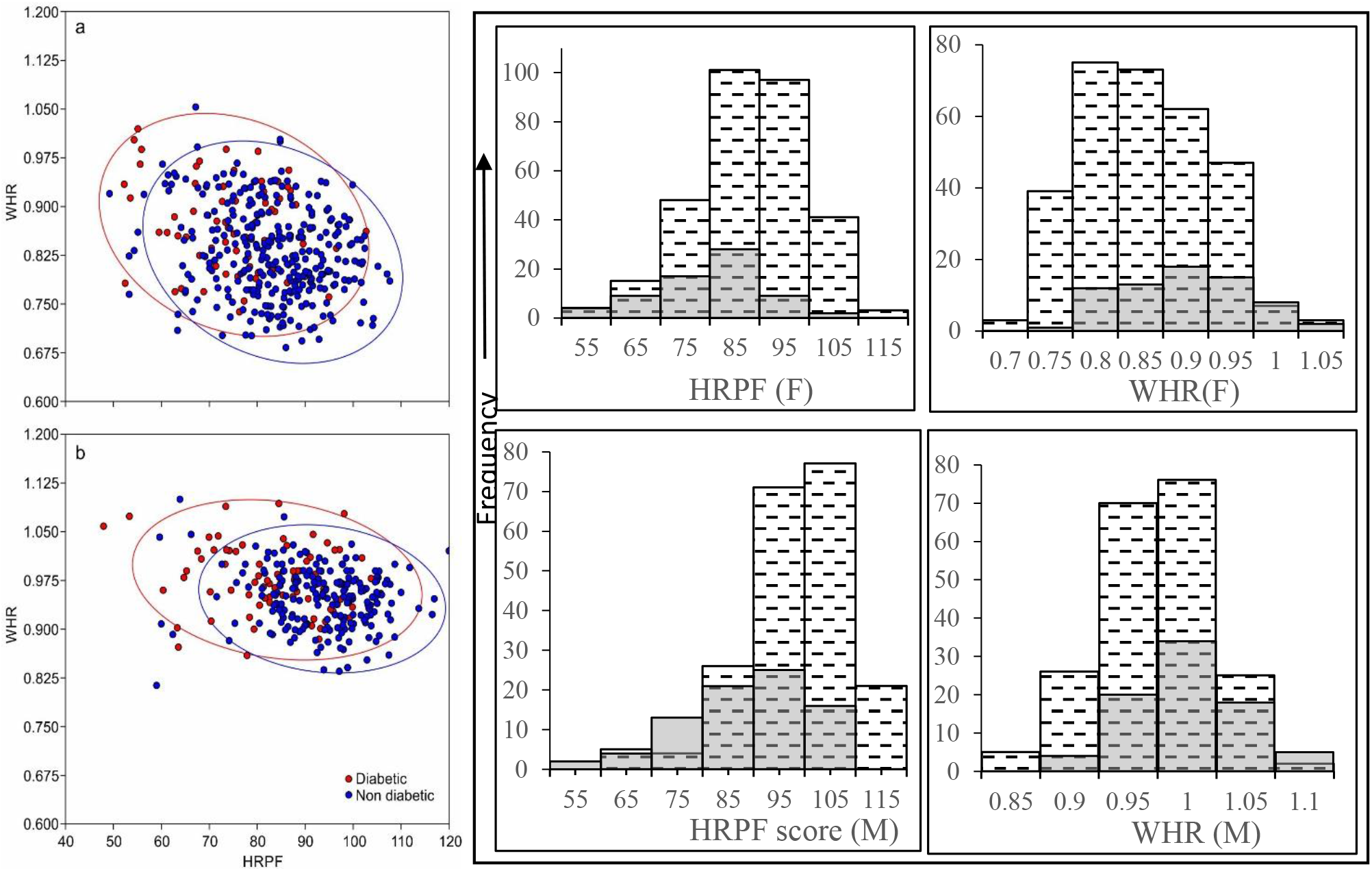
The distributions of diabetics (orange circle) and non-diabetics (blue circle) with respect to HRPF score and WHR (a) females (b) males. The bar diagrams represent the distributions along one dimension at a time (grey bars diabetic, dashed ones non-diabetic. In both sexes the distribution of diabetics have a downward shift along the HRPF axis and an upward shift along the WHR axis.

Since WHR and HRPF are independent predictors of T2DM, one would expect maximum risk for individuals with bad fitness score as well as bad WHR. Simultaneously it would be enlightening to see the risk for individuals that have good functional fitness but bad WHR, or good WHR but bad functional fitness. Since the correlation between WHR and HRPF was weak, it was possible to find individuals in the worst (forth) quartile of WHR but the best (forth) quartile of HRPF and vice versa. When the proportion of diabetics in the four combinations were plotted (figure 5) the maximum incidence (32 out of 59) was in the lowest HRPF and highest WHR class as expected, and lowest (only 2 out of 44) in the diametrically opposite combination. This indicates that the effects of WHR and HRPF are synergistic. However it can be seen that the gradient across HRPF was much sharper than the gradient across WHR. Furthermore in the combination where WHR is expected to have a prodiabetic and HRPF a protective effect, the incidence was only 1 out of 18, but when HRPF was predictive of diabetes and WHR protective it was 12 out of 31. So when cases in which the two predictors were in opposite direction, HRPF was a better quantitative predictor than WHR.

**Figure 5:**
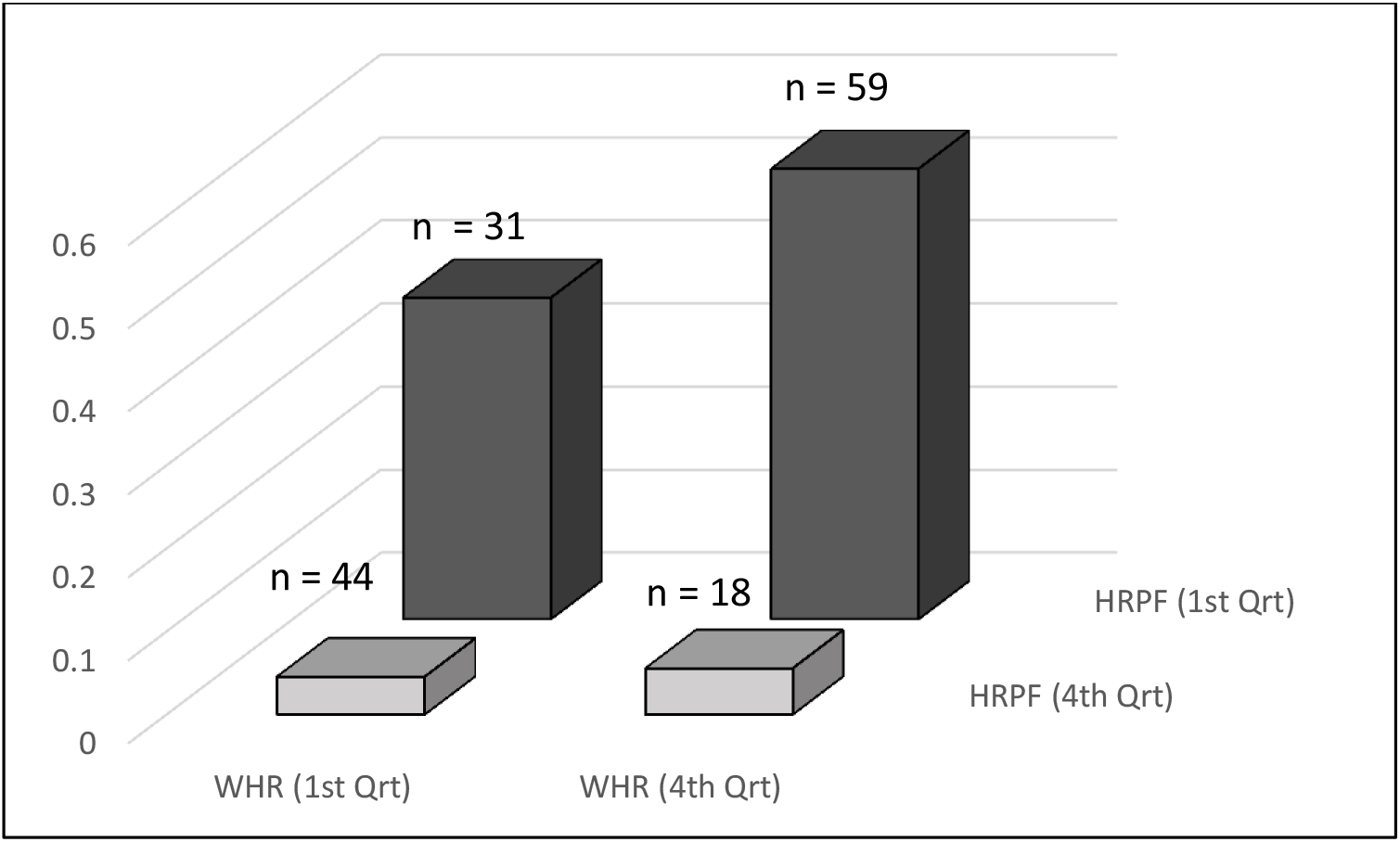
The proportion of diabetic people along the first and forth quartiles of WHR and HRPF. As expected, the proportion in the lowest HRPF and highest WHR quartiles is the largest and highest HRPF and lowest WHR quartiles the smallest. Particularly notable pattern is that individuals that has good HRPF scores had low incidence even when WHR was bad. On the other hand when WHR was good but HRPF bad, the incidence was substantially higher. Pooled data on both sexes where ranking is done separately in males and females.

In order to examine whether the association between lower HRPF scores and T2DM is dominated by the weaker components of fitness or the stronger ones, we ranked all individuals according to each of the component fitness scores and then for each individual totalled the lowermost and uppermost three ranks. The index thus obtained for the weaker dimensions of fitness was a much better predictor of T2DM, the odds ratio for males being OR =13.4 (4.84, 37.14) and females OR = 19.51 (4.47, 84.3). On the other hand the index for the three strongest components had a relatively lower predictive power, OR for males being 8.52 (3.06-23.76); females 4.64 (2.06, 10.4)). The difference between ORs of weakest and strongest components was statistically significant for females (p one tailed 0.039), for males it was not statistically significant but the direction of change was similar. This suggests that weakest fitness dimensions appear to influence the association more than the strongest fitness dimensions for any individual.

Since we observed from the data that there was a significant association between low fitness score and T2DM, we tried to address the question whether diabetes leads to progressive decline of fitness or whether loss of fitness is a predisposing factor for diabetes. The age related decline in HRPF is faster in diabetics than in non-diabetics in males but not in females as shown earlier. If diabetes leads to progressive loss of fitness, we would expect people with longer duration of diabetes to have lower fitness scores. Among the individuals in the sample data on duration of diabetes was available for 52 males and 38 females. In this subsample, we did find a significant negative correlation between HRPF and duration of diabetes in males (r = −0.38, p = 0.01). After correcting for age the correlation was lost. On the contrary, the age-HRPF negative correlation was not lost after correcting for duration of diabetes (corrected r = 0.569, p = 0.01). This suggests that the apparent negative correlation of HRPF with duration of diabetes is likely to be contributed by age alone. In females, on the other hand, there wasn’t a significant correlation between HRPF and duration of diabetes (r = −0.237, p > 0.05). Therefore the prediction of the hypothesis that diabetes leads to a progressive loss of fitness is not supported by data. Although the causal relation cannot be ascertained from this analysis alone, its inability to support progressive effect of diabetes on fitness makes it more likely that loss of fitness predisposes to diabetes.

## Discussion

The different dimensions of functional fitness such as balance, endurance, flexibility, nerve-muscle coordination, muscle strength, core strength and agility captured by component fitness scores were correlated positively with each other but the correlations were weak. This means that the different components of fitness are unlikely to be represented well by a single test or measurement. Fitness is necessarily a multidimensional concept and only morphometry or a single task performance does not reflect on overall fitness sufficiently well. In particular, BMI and WHR were poorly correlated with the functional fitness components and therefore are unlikely to be good surrogates of overall fitness.

The most important finding of the study is that the multidimensional fitness score is a substantially better predictor of T2DM as compared to BMI and WHR in a cross sectional sample. Our study was retrospective and was not a randomized sample from the population. Nevertheless there is no apparent reason why a self-selection bias would lead to lower fitness scores in diabetics as compared to non-diabetics. The study is certainly indicative and should be followed up with prospective studies with better sampling design.

The question whether any particular single component of fitness is a better predictor of T2DM needs to be kept open since we did not find a significant difference in their predictive ability in terms of OR. Nevertheless almost all single components had ORs smaller than the total score which indicates the importance of multidimensional functional fitness score. If some component is found to be consistently the best predictor across different populations, it may become a simple and single useful test in future, but from our sample it appears that it is necessary to look at different dimensions of fitness.

The exercise of taking the lowermost and uppermost scores suggests, on the other hand, that rather than a single component, an individual’s weakest fitness components appear to determine the risk of being diabetic. This means that a balanced and all round fitness needs to be emphasized and only being normal weight or strong in one or two components may not be sufficient. Different types of exercises have differential emphases on particular fitness components. Therefore rather than one particular type of monotonous exercise, diversity of exercises strengthening different components of functional fitness is likely to be a more successful preventive measure. This is an interesting possibility raised by this study which needs more research to explore its translational importance.

The study exposes the extremely limited role of obesity parameters in T2DM either as causal or simply correlated variables. Particularly since south Asia is known to have a substantial number of normal weight type 2 diabetics, it is necessary to move the focus from obesity to better correlated, and if possible demonstrably causal factors. Sarcopenia is a known predisposing factor for insulin resistance (20)(21) which would be reflected in some components of the fitness tests. However in our study balance, endurance and nerve muscle coordination were also good predictors of T2DM without necessarily being very highly correlated with muscle strength. Therefore it is possible that apart from fat and muscle mass, a number of other functional fitness parameters play a role.

The mainstream thinking about the aetiology of type 2 diabetes has undergone many radical changes in the history. Until the 1960s many researchers believed that T2DM had a Mendelian genetic origin (22) and genes predisposing to T2DM contributed to obesity by the tendency to produce large amounts of insulin (22). In this thinking high insulin response was assumed to be causal to obesity. In the following decades the thinking was turned upside down and obesity was held primary and was believed to induce insulin resistance (23–24). The mainstream thinking of over three decades that obesity leads to insulin resistance and further to compensatory hyperinsulinemia is challenged today again by the claims that hyperinsulinemia precedes insulin resistance (2–5) and that high insulin levels are the primary contributors to obesity. While the causal relationship between insulin, insulin resistance and obesity is being debated, meta-analysis of the obesity insulin resistance showed that the correlation itself is quite week, explaining less than 15 % variance in most studies (9). Independent of the debate on the direction of the causal arrow, obesity-insulin resistance relationship by itself is inadequate to account for type 2 diabetes. Therefore there is an increasing need to look beyond obesity and look for stronger causal or at least better correlated factors.

At the same time it is necessary to elaborate on the possible theoretical underpinnings of the association between physical fitness and T2DM. In evolutionary medicine a number of hypotheses for the origin of T2DM have been suggested. The classical thrifty gene (23) and thrifty phenotype (25) hypotheses are obesity centred. There are alternative hypotheses based on behavioural and reproductive strategies or life-history strategies (26–28) which are not obesity centred, although they might allow a correlation with obesity. In these hypotheses, physical strength, social ranking and thereby reproductive opportunities play a more important role than obesity. A definite role of brain and neuronal circuits is being highlighted by a number of recent studies (29–31). In an ancestral environment, physical fitness is expected to play a role in deciding behavioural and reproductive strategies and therefore a change in metabolic and neuroendocrine make up may follow loss of physical fitness. For example, a physically weak individual is most likely to be a subordinate individual in a primate social hierarchy and accordingly needs to change its foraging, social and reproductive strategies (32–33). The position in social hierarchy might be lost by the loss of one or more components of fitness. Here the weaker components are likely to matter more than the stronger components. The strategic changes required on being weak are accompanied by metabolic, endocrine and immunological fine tuning (31–34). Although the relationship between physical fitness and social hierarchy has changed in the modern human society, human physiology is still likely to be responding according to the evolved ancestral optimization. This is the potential theoretical underpinning of the significant association between functional fitness and T2DM.

The BILD clinic from where we took data for the retrospective study, had established a scoring system for assessing the different dimensions of functional fitness for a different purpose. We used the same scores in our analysis although they were not primarily optimized for the hypothesis under investigation. It is possible with more focussed research that simpler but better optimized tasks for multidimensional functional fitness can be designed specifically for more reliable prediction of endocrine and metabolic disorders. If our findings are found consistent and reproducible in other studies, simpler but better predictors can be devised to find useful clinical applicability.

## Methods

### Statistics used

We used Pearson’s correlation analysis to examine the interrelationship between different fitness components and their relationship with age and obesity parameters. Bonferroni correction was applied to use a conservative significance level since multiple correlations were being performed. Since T2DM is an age related disorder and since age had a significant negative correlation with fitness, age corrected fitness scores were used for further analysis.

The male and female populations were divided into quartiles according to HRPF, individual fitness components, BMI and WHR. Odds of being a diabetic were calculated for each of the quartiles. Odds ratio (OR) between the 1st and 4th quartiles was used as a predictor. For the functional fitness components the odds of being diabetic were expected to be higher in the lowest quartile therefore the ratio of first to forth quartile was expressed. In the case of BMI and WHR, odds in the forth quartile were expected to be higher and therefore the ratio was expressed as forth to first quartile. The significance of difference between odds ratios was tested by z transformation of log OR difference using the standard errors of OR. We also looked at the odds ratios in the total population where males and females were ranked and quartiles identified separately and then pooled together.

To examine whether the best or the worst fitness components contributed more to the association between HRPF and T2DM, we ranked individuals for each functional fitness component independently and then added the 3 highest and 3 lowest ranking scores of every individual and looked at odds of being diabetic across their quartiles.

Multivariate normality was tested using (35) omnibus. Since the data were not normal for both females (Ep = 15.4, p < 0.004) and males (Ep = 20.82, p < 0.001), we used non-parametric multivariate analysis of variance or PerMANOVA (36) with 9999 permutations to test the null hypothesis that the centroids and dispersion of the groups, in this case diabetic and non-diabetic, as defined by measure space are equivalent for all groups. PerMANOVA was performed using PAST 3.22 (37)

## Author contributions

PP and MW conceptualized the study. MW, PP and PL designed and standardized the protocol. HV and MW did the statistical analysis and inference part of the study. SK, DB and MJ contributed to the standardization of exercise protocol part. MR, NB, MN and SB took care of exercises/tasks performed by participants.

Authors declare there is no conflict of interests.

## Acknowledgments

We than the management of Deenanath Mangeshkar Hospital and Research Center that supports the BILD clinic, Mr. Deepak Desai and Mrs. Sarita Desai for financing the study; Dr. Charu Bapaye, Mrs. Poorva Joshi, Mrs Swati Mantri and management of Kamayani School for their support in the conceptualization stage, Mrs. Prajakta Gurjar, Dr. Meeta Peer (Mr. Nanasaheb Bhanage Rehabilitation Center) for administrative support.

