## Supplementary Information for "A multidimensional functional fitness score is a stronger predictor of type 2 diabetes than obesity parameters in cross sectional data"

Short title: Fitness score based prediction of type 2 diabetes

Pramod Patil[2] , Harshada B Vidwans [1],Poortata S Lalwani[3], Shubhankar A Kulkarni[1] Deepika Bais[1],Manawa M Diwekar-Joshi[1], Mayur Rasal[2], Nikhila Bhasme[2], Mrinmayee Naik[2], Shweta Batwal[2], *Milind G Watve[1]

1 Indian Institute of Science Education and Research, Pune (IISER – P) India.
2. Deenanath Mageshkar Hospital, Pune India.
3. Department of Psychology, University of Michigan, Ann arbor, MI - 48109, USA

Corresponding author: Milind G Watve,
Indian Institute of Science Education and Research, Pune (IISER – P) India.
Dr. Homi Bhabha Rd, Ward No. 8, NCL Colony,
Pashan, Pune, Maharashtra 411008

Phone(O) : +91- 020- 25908093
Fax (O) : +91-020-25899790

Supplementary information:

**Health related physical fitness (HRPF) assessment protocol**

BILD (Behavioural Intervention for Life style Disorders) clinic has standardized a set of 15 task performance tests. Many of these tests are adopted from published literature with or without modifications to suit the age group, local culture and to the specific sequence of tasks to perform. Some of the tasks are newly conceived and standardized. Every individual is asked to perform the tasks and is observed while performing the tasks. Each task performance is scored according a set of task specific norms. The total score reflects a sum of different components of fitness. The 15 tasks together assess 8 components of fitness namely abdominal plasticity, balance, endurance, flexibility, nerve-muscle coordination, muscle strength, core strength and agility. Participants get a total score out of 100, each fitness component is separately scored out of 10 or 15. The task wise protocols are as follows:

1. Morphometry and abdominal plasticity: This comprises of Body mass index (BMI), waist hip ratio (WHR) and abdominal plasticity (AP).
   a) BMI and WHR was calculated according to standard protocols (1,2).
   b) Abdominal plasticity: AP is the ratio of waist circumference in the inflated and deflated state. AP is expected to reflect visceral fat and abdominal muscle tone.
      Waist circumference was measured by wrapping tape around the top of the hip bone (illiac crest) at the level of umbilicus. The mean of two consecutive measurements that do not differ by more than 1 cm was recorded.
      Waist deflated: In a standing position the participant is asked to deflate his/her belly by sucking/ retracting it inside as much as possible.
      Waist inflated: In a standing position the participant is asked to inflate his/her belly as much as possible.
      Abdominal plasticity score was calculated as 20 x (inflated circumference – deflated circumference)/ (deflated waist circumference)*75. The score generally ranges between 0 and 15. We did not find any individual exceeding a score of 15.
2. Balance: Three types of balance tasks are performed, each carries five marks. Total 15 marks for balance.
   a) Stork balance stand test (3): Participant is asked to stand straight upright with both hands rested on the hips then asked to position one foot against the inside knee/thigh of the standing leg and stand stably for 30 seconds. As the second grade task they are asked to raise and join their hands above the head and stand stably for 30 seconds. As the third grade task they are asked to stand in the position of grade 2 with eyes closed. The participants are first given 30 sec to practice the balance before recording score. Each grade was started as a new attempt, without continuing from the previous grade, this was done in order to avoid fatigue of the standing leg. Participants were free to choose between left or right standing leg. A score of two was given to each successful grade completion. Grade 3 had a high level of difficulty and less than 2 % participants completed it. Therefore although the test was finally counted as out of 5, exceptional individuals could obtain one bonus point by completing grade 3. Score was obtained by multiplying with 2.

b) Dynamic balance (4): In this test participant is asked to maintain single-leg balance while raising the contralateral leg high in four different directions one after the other and holding it for 2 seconds each. The leg is not allowed to rest between any of the four positions. All actions are to be completed without losing balance or touching the ground. Four positions were achieved in following sequence: forward, sideways, back and knee high position. For accurately performing the entire test a full score of 5 is given. For every position not completed 1 mark is deducted while for loss of balance but regained without touching the ground a 0.5 mark is deducted. For touching the ground between transitions one mark each is deducted.

c) Wear and remove socks: In this test participant stand upright on a firm hard ground with no socks or footwear. In the upright position, they are asked to raise one knee and wear and remove socks without bending forward or tilting the leg or taking any support. Best attempt out of three is considered for scoring. For smooth and flawless act 5 marks are given and for every compromise such as bending, losing balance and touching the ground or tilting the leg 1 mark each is- deducted. For losing balance but regaining without touching the ground 0.5 mark is deducted.

3. Endurance: There are two test for endurance each carries 5 marks. Total 10 marks.

a) Breath holding time is used to test respiratory endurance. Participant is asked to breathe in slowly as deep as possible using accessory muscles of respiration and hold the breath in inspired state for as much time he/she could. Once the breaking point is reached participant breathed out slowly. Three such consecutive breaths are taken without gap. This entire duration of three breathes is measured in seconds. The score was calculated by dividing the average number of seconds by 10. Since the maximum observed value was not exceeding 50secs.

b) Arm curl test for muscle endurance (5): The participant is asked to perform as many arm curls as possible in 30 seconds with a weight of 5 Kg for males and 3 Kg for females. The score was calculated by diving the total number of arm curls by 5, since the maximum observed was not more than 25.

4. Flexibility: (6)(7) Each test carries 5 marks, total 10 marks for flexibility.

a) Sit and Reach/ Trunk Flexion Test: In this test participant is asked to sit on the floor with legs stretched out straight ahead. The soles of the feet are placed flat against the box. With the palms facing downwards, and the hands on top of each other or side by side, the participant reaches forward along the measuring line as far as possible. The most distant point (cm or in) reached with the fingertips is noted in centimetres. The score is calculated as distance divided by 9. Maximum score is 5 marks.

b) Sit and split: Participant is asked to split on the ground facing a wall with feet touching the wall. To support the body participant holds the rope attached to the wall in front and pulls his/her body forward against the rope and spreads his/her legs apart and tries to go as close as possible to the wall. The distance between the two heals is measured in cm. The score is calculated as if the distance is less than 90 then score is zero otherwise 5*(distance- 90)/ (2* (leg length – 90) out of 5.

5. Nerve-muscle coordination: For testing the nerve- muscle coordination alternate hand wall toss test was done. A mark is placed at a distance of two meters from the wall.

Participant is asked to stands just behind this line facing the wall and throws a ball from one hand in an underarm action against the wall, the ball bounces from the wall to the ground and then participant attempts to catch it with the other hand. The ball is thrown back against the wall by the hand in which ball was caught and then the initial hand tries to catch it. Total time given is 30 sec. and maximum number of successful attempts are counted. If a ball is missed the trainer provides another without delay. Participant is allowed to practice it for 30 secs before the actual test. Score is calculated as the total number of catches divided by 4. Maximum number of marks is 5.

6. Muscle strength: There are three test for measuring muscle strength, static handgrip test, broad jump test carries 5 marks each and sitting –rising test is carrying 10 marks. Total 20 marks for muscle strength.
   a) Static handgrip strength test (8): The participant is asked to hold the dynamometer in the dominant hand. Arm should lie by the side of the body vertically down. The handle of the dynamometer is adjusted if required - the base should rest on first metacarpal (heel of palm), while the handle should rest on middle of four fingers. When ready the participant should press the dynamometer with single maximum isometric effort. No other body movement is allowed. Only dominant hand is tested with total three attempts. The best of all three readings is considered. Score calculated as for males the reading was divided by 8 and in females divided by 6. Score was given out of 5.
   b) Standing broad jump test: The participant is asked to stand behind a line marked on the mat with feet slightly apart. A two foot take-off and landing is used, with swinging of the arms and bending of the knees to provide forward drive. The participant attempts to jump as far as possible, landing on both feet without falling backwards. Three attempts are allowed. The measurement is taken from take-off line to the nearest point of contact on the landing (back of the heels). The longest distance jumped out of three attempts is considered. The score is calculated by distance multiplied by divided by height of the individual. The score is given out of 5.
   c) Sitting-rising test (9): Participant is asked to stand upright on a firm plain ground with feet at shoulder distance apart. Without worrying about the speed of movement, participant has to try to sit cross-legged and then to rise from the floor, using the minimum support that is needed. It is very helpful to keep both hands extended forward while performing the test. The best is to use no support while sitting and standing. Marking system for sitting- A half mark are deducted for loss of balance. 1 mark is deducted for every support taken. Marking system for rising- A half mark is deducted for loss of balance. 1 mark is deducted for every limb support taken. Subject is given 3 attempts and scored out of 5 for sitting and out of 5 for standing, thus, out of a total of 10. Maximum marks that can be deducted out of five are 4.
7. Agility: There are two test for agility shuttle run test carries 5 marks and complex new motor skill learning test carries 10 marks, total 15 marks.
   a) Shuttle run test. On the signal "ready", the participant is asked to place their front foot behind the starting line. On the signal, "go!" the participant sprints to the opposite line, picks up a cone, runs back and places it on or beyond the starting line. This is repeated at a stretch till all four cones are shifted and total four trails

are complete. Participant is given rest after this task before proceeding to the next task. The time to complete the test in seconds to the nearest one decimal place is recorded. The score is calculated as (30 – time)/4. Score was given out of 5.

b) Complex new motor skill learning:
Ladder drill test: Participant is asked to stand at one end of the ladder on one side, facing rest of the ladder length. To start with participant should stand such that ladder is on his/her left side. In this position, participant has to move right leg ahead (step 1) and place it parallel and outside of the first bracket. Then place left foot inside of the same bracket (step 2). Then the right leg again takes one step at the same position (step 3). Then left foot is taken out of the bracket and is placed along the side of the same bracket in between right foot and bracket. First the sequence of movement expected is demonstrated in a slow purposeful manner. Participant is given one trial (half of the ladder) to understand and practice the sequence. The total time taken to complete sequence accurately along the length of the ladder is noted. Ladder should not be touched and balance should not be lost. The task is to be completed as fast as possible and participant is motivated to perform better with each trial. To be done three times (start to end, end to start and again start to end direction) on the same side of ladder. Time for each attempt is noted. The least time taken is scored. Time taken to finish the drill from one end to the other for three individual attempts is recorded. The score is calculated as, if time is more than 30 then zero otherwise (30 – time)/ 2.5. Score was out of 10.

8. Core strength: There are two tests each carries 5 marks, total 10 marks for core strength.

a) Upper abdominal strength test (UAST): The participant is asked to take supine position. Legs flexed at knee and hip joint at a convenient angle. In all grades participant should perform minimum 5 repetitions in 30 seconds. Grade 1: Participant is asked to hold the cotton belt in both hands and rounds it below both feet and is asked to perform curl ups by pulling the body with the help of belt. Grade 2- Abdominal curls performed without the support of the belt. Participant outstretched both hands in front. He/she can use hands for getting thrust- forward motion to lift the body. Grade 3- Abdominal curls performed with hands crossed over chest. Grade 4- Abdominal curls performed with both palms placed behind the head with elbows facing forward. Grade 5- Abdominal curls performed with palms placed behind the head and elbows in the plane of the head. 5 repetitions in 30 seconds were required to complete each grade. Unable to perform even grade 1 is considered as Grade 0. Successfully completed grade is considered (0 to 5). Maximum score is 5.

b) Lower abdominal strength test (LAST): In supine position participant is asked to lift both the legs from hips without flexing in knees. Time was measured for which a particular position has been achieved. Every position is to be maintained for minimum of 10 seconds. Grade 5- Performing straight leg raise (SLR) upto 15 degrees. Grade 4 - Performing SLR upto 30 degrees. Grade 3- Performing SLR upto 45 degrees. Grade 2- Performing SLR upto 60 degrees. Grade 1- Performing SLR up to 90 degrees. If grade 3 performed successfully then go to grade 4 and then 5. If grade 3 is not done then shift to grade 2 and if not then grade 1. Failure to perform grade 1 is considered as grade 0. Successfully completed grade was mentioned. (0 to 5). Score is given out of 5.
